# Temporal persistence and structural organization of neuronal avalanche dynamics

**DOI:** 10.64898/2026.08.10.743923

**Authors:** Gianmarco Cafaro, Marianna Angiolelli, Matteo Demuru, Gabriele Casagrande, Emahnuel Troisi Lopez, Mario Quarantelli, Carmine Granata, Damien Depannemaecker, Gian Marco Duma, Silvia Scarpetta, Pierpaolo Sorrentino

**Affiliations:** Department of Physics “E.R. Caianiello”, University of Salerno, Fisciano, Italy; Epilepsy and Clinical Neurophysiology Unit, IRCCS Eugenio Medea, Conegliano, Italy; Department of Motor Sciences and Wellness, University of Naples “Parthenope”, Naples, Italy; Aix-Marseille Univ, INSERM, INS, Institut de Neurosciences des Systèmes, Marseille, France; Department of Education and Sport Sciences, Pegaso University, Naples, Italy; Institute of Biostructures and Bioimaging, Italian National Research Council, Naples, Italy; Institute of Applied Sciences and Intelligent Systems of National Research Council, Pozzuoli, Italy

## Abstract

Brain activity can be understood as a sequence of neuronal avalanches, i.e., transient episodes of coordinated activation that emerge across scales, from individual neurons and local networks to whole-brain dynamics. Avalanches are typically characterized by features such as size, duration, number of active components, and the silent time separating consecutive events. Although these features have been extensively characterized through their marginal distributions, their temporal organization and dependence on the underlying brain architecture remain poorly understood, leaving us without a framework for embedding neuronal avalanches within slower brain dynamics.

Here, we analyzed eyes-closed resting-state magnetoencephalography recordings and the corresponding structural connectomes from 30 healthy participants to investigate the dynamics of avalanche sizes and silent times. We found that large avalanches preferentially followed short silent times, whereas small avalanches were more likely to occur after long silent periods. Based on the empirical joint distributions of avalanche size and silent time, we could define four types of events occurring above chance levels (avalanche large or small, preceding pause long or short). Mixed categories—combining a small value of one feature with a large value of the other—occurred more frequently than expected, while same-category events happened less often than chance. Furthermore, consecutive events tended to remain in the same category, a phenomenon referred to as persistence. We next investigated whether a brain region’s connectivity profile shapes its propensity to participate in avalanches of different sizes. More strongly connected regions participated most often in small avalanches, whereas weakly connected regions were preferentially recruited during large avalanches. This pattern may reflect the greater sensitivity of highly connected hubs to fluctuations propagating through the network, resulting in frequent but spatially contained events. By contrast, the recruitment of more peripheral regions may require broader and stronger collective activity, occurring only during rarer, large-scale avalanches. In contrast, regional participation showed no clear association with the silent time preceding an avalanche. Together, these findings show that neuronal avalanches are neither temporally independent nor anatomically unconstrained: their sequence retains a memory of preceding events, while structural topology shapes which regions are recruited as avalanches grow. By connecting avalanche dynamics with slower temporal organization and the structural connectome, our results provide a multiscale framework for understanding how transient events are embedded within ongoing brain activity.

## 2 Introduction

Neuronal avalanches are a widely studied framework for investigating collective neural dynamics across multiple spatial scales, ranging from local neuronal populations to whole-brain activity. Originally identified in organotypic cortical slice cultures [1] and dissociated cultures [2, 3], and later observed in-vivo in invasive and non-invasive recordings [4–8], neuronal avalanches are characterized by scale-invariant distributions of size and duration, a hallmark often associated with collective dynamics operating near criticality [8–10]. Near-critical dynamical regimes are posited to support efficient information transmission, large dynamic range, and optimal information processing in neural systems [11–15].

While the statistical properties of avalanche size and duration have been extensively investigated, how these are nested within concurrent, slower dynamics remains unclear. In particular, the intervals separating consecutive avalanches, commonly referred to as silent times or inter-avalanche intervals, provide important information about the slow mechanisms governing the inter-avalanche dynamics. Silent time statistics may provide information complementary to avalanche size and duration distributions. For example, experiments on granular systems and numerical simulations of emulsions have shown that the same probability distributions of avalanche size and duration can exhibit different silent time distributions [16]. In systems composed of independent events, silent times are expected to follow simple exponential statistics. Deviations from this behavior indicate the presence of temporal correlations and suggest that consecutive events are not generated independently. In neural systems, silent time distributions exhibit a complex structure characterized by multiple scaling regimes and long-range temporal correlations [17–20].

Recent work has shown that the temporal succession of neuronal avalanches displays non-trivial dependencies between consecutive events. The size difference between an avalanche and the preceding one, referred to as amplification when positive and attenuation when negative, depends on the silent time separating the two events. This relationship with the size propagation is further modulated by ongoing alpha oscillations, revealing a coupling between avalanche propagation and ongoing oscillatory brain activities [21]. These findings suggest that neuronal avalanches are not isolated events but are influenced by longer-timescale dynamical processes that shape their rhythms.

Interestingly, similar temporal organization has recently been reproduced in neuronal cultures and in a Kuramoto-like model incorporating excitatory and inhibitory interactions, showing that scale-free avalanches, synchronization, and non-trivial temporal correlations can emerge from the collective dynamics of coupled neuronal populations [20]. Together, these experimental and theoretical findings suggest that the dynamics of avalanches cannot be understood as a sequence of independent cascades but should instead be regarded as a collective process shaped by history and guided by the evolving synchronization state and excitatory–inhibitory interactions of the neuronal network.

Previous work has shown that avalanche size is related to the preceding silent interval and that the interval between consecutive avalanches is associated with whether subsequent activity is amplified or attenuated. These findings indicate that the timing and propagation of neuronal avalanches are interdependent. Such local dependencies may, however, represent the observable expression of a slower dynamical organization spanning multiple events.

This possibility is consistent with the broader view of spontaneous brain activity as a succession of metastable network states. Experimental studies have long documented spontaneous alternations between periods of enhanced and reduced excitability, commonly described as up and down states [22–24]. Theoretical work has likewise shown that collective network dynamics can generate multiple attractor-like states that persist substantially longer than individual activity cascades, thereby organizing activity over extended timescales [25]. Within this framework, individual avalanches need not constitute independent events, but may instead represent transient expressions of an underlying dynamical state. Indeed, avalanches have been proposed to emerge near the transition between the replay and non-replay of spatiotemporal activity patterns, suggesting that they may correspond to fragments of longer-lasting network dynamics [26–28].

Based on these observations, we hypothesized that neuronal avalanches are organized into persistent dynamical regimes. Specifically, if such regimes exist, particular combinations of avalanche size and silent interval should recur in structured sequences, producing temporal correlations that cannot be explained solely by interactions between consecutive avalanches. We further reasoned that the expression of these regimes should be constrained by the brain’s structural architecture. Empirical data showed that the spatio-temporal spreading of neuronal avalanches is constrained by the underlying structural connectome [29, 30]. Accordingly, whole-brain modeling studies have shown that the correspondence between structural connectivity and functional dynamics is maximal near criticality, indicating that collective activity patterns are strongly shaped by anatomical connectivity [27]. Hence, the topological position of individual regions within the structural scaffolding may determine how they are recruited across avalanches with different size and timing characteristics.

To test whether neuronal avalanches are organized into persistent dynamical regimes and whether this organization is constrained by the structural connectome, we jointly analyzed avalanche size and the silent interval preceding each event. We first quantified the statistical dependence between these variables and used their joint distribution to define distinct avalanche states. We then examined whether these states persisted across successive events, as expected if the underlying network retains a dynamical memory spanning multiple avalanches. Finally, we characterized the spatial recruitment patterns associated with these states and assessed whether regional participation was shaped by the topology of the structural connectome. Our findings reveal that avalanche dynamics exhibit persistent temporal organization and that the recruitment of brain regions across events of different sizes is constrained by the underlying structural architecture.

## 3 Results

Initially developed to characterize critical dynamics in spiking neural networks, neuronal avalanches have since been extended to the study of whole-brain activity using both invasive and non-invasive techniques. In particular, in EEG and MEG recordings, continuous signals can be discretized by considering deviations of each source signal from its mean value exceeding a given threshold, typically around 3 standard deviations, where the distribution of fluctuations starts to depart from Gaussian behavior. By grouping consecutive discretized events across all sources, it is possible to identify bursts of activation (avalanches), whose duration and size—defined as the number of discrete events occurring across all active regions—have been shown to follow power-law distributions. The observed scale-free behavior of these quantities, together with other signatures of criticality, suggests that brain activity may self-organize near a critical regime, at the boundary between quiescence and a self-sustained activity regime [13] [31].

### 3.1 Size - Silent Time Joint Probability Distribution

Although the relationship between avalanche duration and size is well established and has been extensively characterized in both empirical data and theoretical models, how avalanche size depends on the time elapsed since the preceding event remains an open question [16, 21].

As a preliminary step, we followed the approach introduced in [5] to estimate the activity threshold used for avalanche detection (Fig.1a). We then quantified the size of each avalanche and the silent interval separating it from the preceding event (Fig.1b).

**Figure 1:**
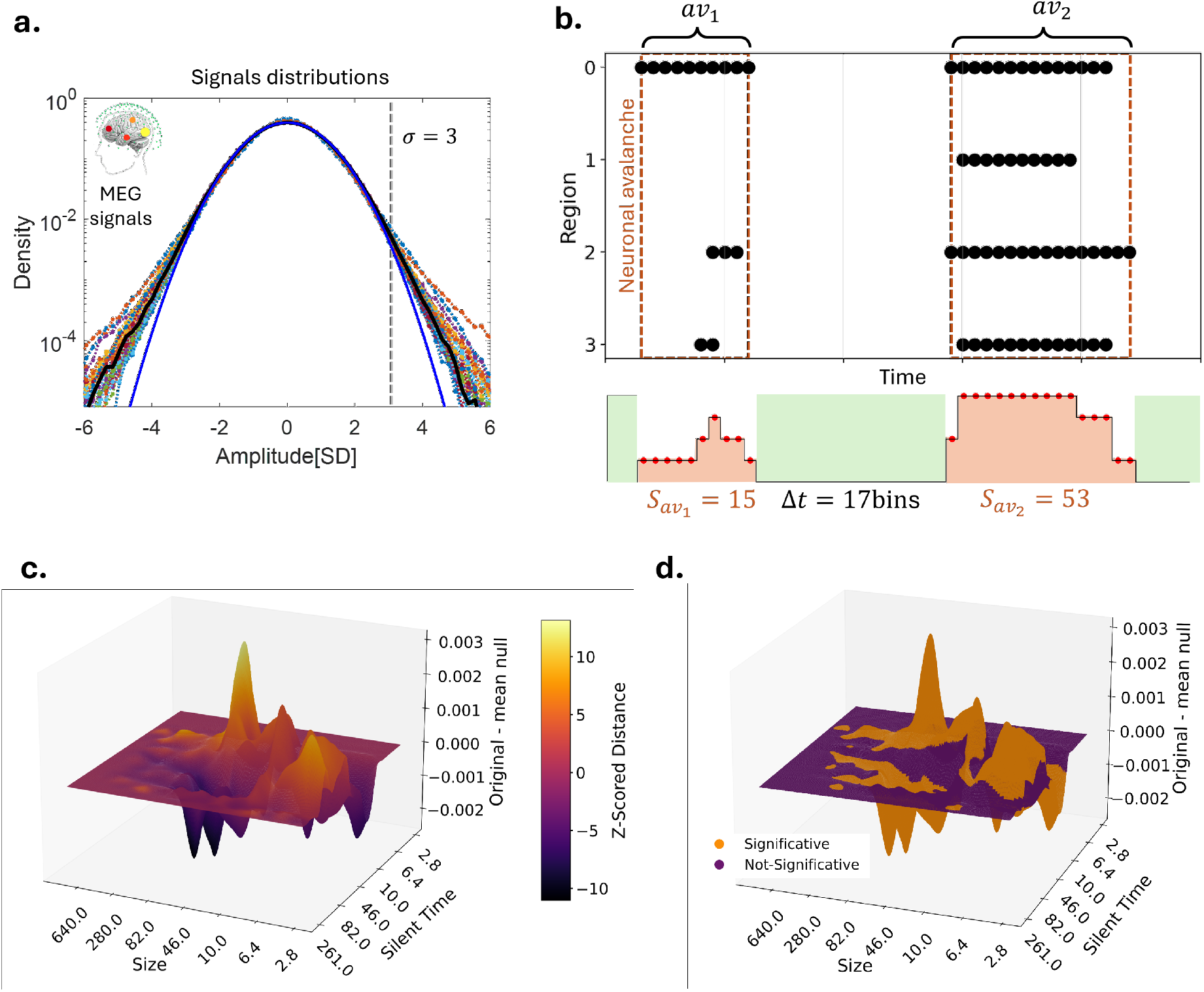
Identification of neuronal avalanches and and Joint distribution of size and silent time. Panel (a,b) Probability density of the Z score normalized MEG signal amplitude. The colored curves in the background are the probability densities for all individual subjects (n = 30 subjects; for each subject we pooled all individual MEG channels). The black curve is the grand average over all subjects. The blue curve is the best Gaussian fit for the grand average. We notice that the empirical probability density starts deviating from the Gaussian fit around *σ* = *±*3*SD*. As usual, the most extreme point in each excursion beyond a threshold h is treated as a discrete event. A representative raster of discrete events across 4 source reconstructed signals is shown in panel b (top). An avalanche is defined as a sequence of temporal windows *ɛ* = 3*ms* with at least one event in any of the regions activity preceded and followed by at least one window with no events in any of the regions (bottom). Avalanche’ size, *S_i_*, is the number of events occurring in the time bins that belong to it. Silent times are the intervals between avalanches. Panel (c) The joint distribution of avalanche’s size and its preceding silent time is computed. Difference between the joint distribution obtained from the original avalanches’ sequences and the average joint distribution derived from the null model, is shown. The plot is colored according to the z-scored difference (Delta), computed as the deviation from the null model mean divided by the standard deviation of the null model distribution. Panel (d): Same as Panel (c), but highlighting only the points where the absolute z-scored distance from the null model (Delta) exceeds 2.

Next, we revisited the relationship reported in [32] using an approach based on the joint probability distribution of the size of the avalanche and the preceding silent time. By quantifying how frequently different combinations of these two variables occurred, we quantify their statistical coupling against a null model in which the temporal relationship between avalanche sizes and silent time intervals was destroyed. This procedure allows us to assess deviations from independence while preserving the marginal distributions of the two quantities.

We identified four regions in the size - silent time space that significantly deviate from the fully random case (Fig.1, panel c). This subdivision, defined by the combination of the boundaries along the two axes, allows us to assess whether a sequence consisting of an avalanche of a given size followed by a silent time of a given duration occurs more frequently than expected by chance. Considering that a z-score larger than 2 corresponds to a probability of approximately 97% of rejecting the null hypothesis, we highlighted the four continuous regions exceeding this threshold Fig.1, panel d).

The results reveal a clear inverse relationship between avalanche size and the preceding silent time: when the time interval between two consecutive avalanches is short, the size of the subsequent avalanche tends to be big, and vice versa. These results naturally partition the size–silent time space into four regions corresponding to combinations of small and large avalanche sizes and short and long silent intervals. In the following sections, we investigate whether this organization is dynamically organized as to give rise to persistent dynamical patterns spanning the duration of multiple avalanches.

### 3.2 Neuronal Avalanche Dynamics Reveals Multiple Dynamical Regimes

To independently assess whether the boundaries emerging from the joint size–silent time organization are also reflected in the marginal distributions of the two variables, we analyzed the distributions of avalanche size and silent time separately using different models. The presence of corresponding boundaries in the marginal distributions would support the interpretation that these states reflect transitions between distinct avalanche-generating regimes, rather than arbitrary partitions of a single continuously varying process.

We first analyzed the probability density functions (PDFs) of avalanche sizes and silent times separately, fitting both distributions above a minimum cutoff value with power laws with exponential cutoffs to account for finite-size effects. For both observables, the truncated power-law model provided a better description of the lognormal. For avalanche sizes, the estimated exponent of the truncated power-law fit is approximately (1.45) Fig. 2, panel a) (left), close to the value predicted by mean-field branching-process theories. For silent times, we obtain an exponent of approximately 1 for truncated power-low. Although these fits provide a satisfactory description of the events above an arbitrary minimum, they do not capture the full shape of the empirical distributions. Indeed, the distributions, especially the size distribution, deviate from the power law at low values, suggesting the possible coexistence of different statistical regimes. Typically, low values are discarded on the account of noise. However, this is typically done without strong evidence.

**Figure 2:**
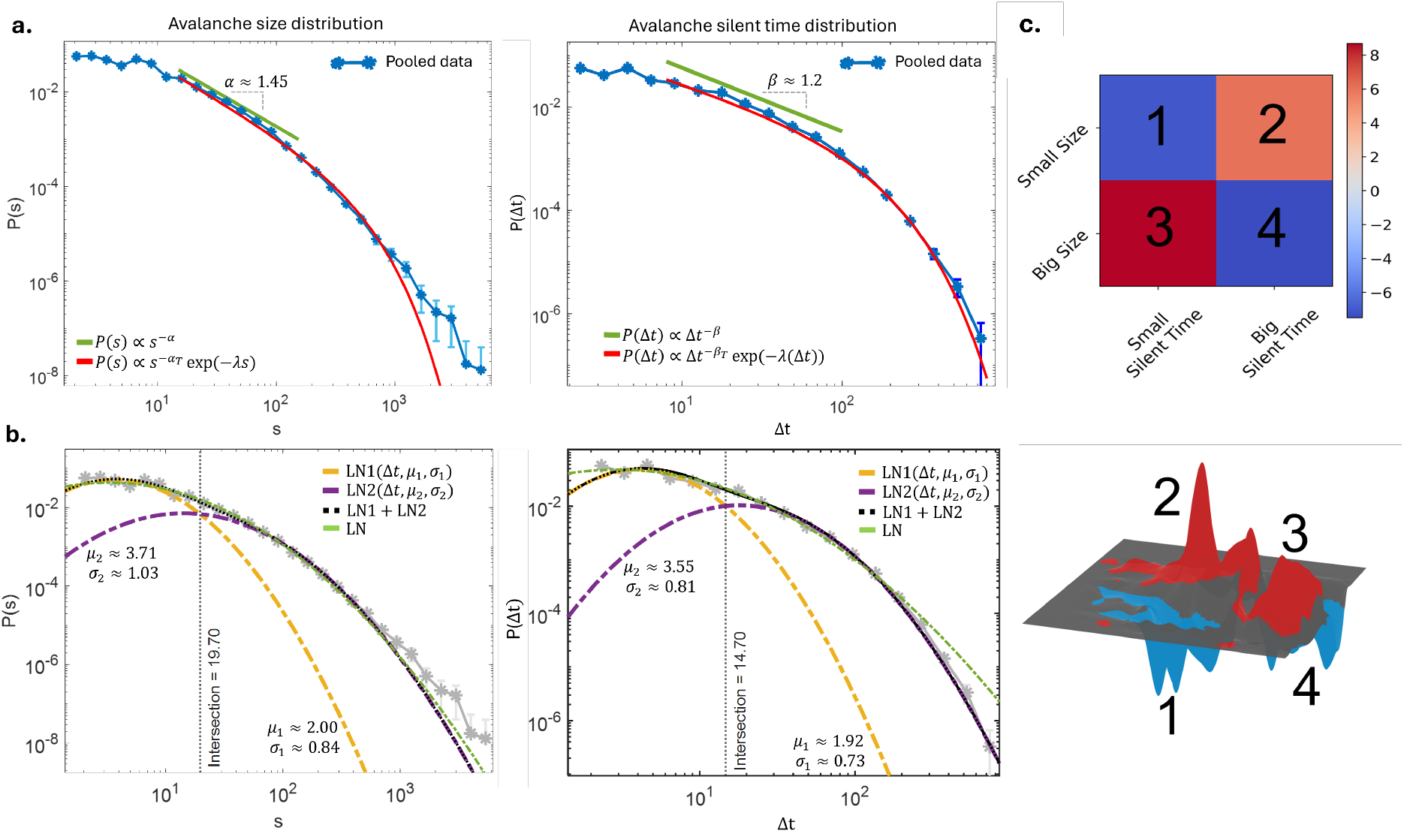
Avalanches as a mixture of different processes. Panel (a): Probability density function (PDF) of avalanche’s size *s* (left) and silent time Δ*t* (right). Blue dots represent the empirical data, while the green and red curves indicate the power-law and truncated power-law fits, respectively. Error bars correspond to twice the estimated standard deviation of the empirical probability in each bin, assuming binomial counting statistics. Specifically, if *P* (*x*) denotes the empirical probability of observing an av<u>alanche size wi</u>thin the interval [*x, x* + Δ*x*], the corresponding uncertainty was estimated as *P* (*x*) = 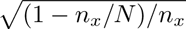, where *n_x_* is the number of avalanches falling in the interval [*x, x* + Δ*x*], and *N* is the total number of avalanches. Error bars correspond to 2*P* (*x*) following [33]. The power-law exponent was estimated using [34]. Restricting the fit to the scaling ranges highlighted by the green lines, yielded exponents of *α ≈* 1.45 and *β ≈* 1.2, respectively. The corresponding Kolmogorov–Smirnov distances were *D* = 0.02 for avalanche size and *D* = 0.03 for silent time. Model comparison showed that, for avalanche size, the power-law distribution was strongly favored over an exponential distribution (log-likelihood ratio *R* = 301.05, *p* = 9.11*e^−^*^11^). In contrast, for silent time, the comparison did not provide significant evidence in favor of either model (*R* = *−*73.56, *p* = 0.13, n.s.). When the upper cutoff *x_max_* was not imposed, a truncated power-law distribution, depicted in red (*α_T_* =1.45, *β_T_* = 1.0) was significantly favored over a log-normal distribution for both avalanche size (*R* = 242.42, *p* = 2.18*e^−^*^16^) and silent time (*R* = 74.20, *p* = 0.006). Panel (b): Probability density function (PDF) interpolated using a mixture of lognormal distributions for size (left) and silent time (right) values. The yellow and violet curves represent the first and second components of the mixed 2-lognormal distribution, respectively. Together, both components generate the overall distribution shown by the black curve. The green curve represents the one-component lognormal distribution. The dotted vertical line indicates the point of intersection between the two components. The green curve represents the single-component lognormal distribution. Grey points correspond to the empirical PDF values. Panel (c) (top):For each avalanche, we compute the probability of belonging to one of the two components of the 2-lognormal mixture distribution. Co-occurrence matrix of the four possible combinations of sizes and pauses, showing the percentage deviation of the empirical probability from the mean probability expected under the null model, as indicated by the color scale. Each cell is assigned an arbitrary number to identify the corresponding combination of mixture components. Panel (c) (bottom): Deviation of the joint probability distribution from the null model. The distribution of sizes and silent times is divided into four regions according to the sign of the deviation from the null expectation. Each region is labeled consistently with the corresponding component combination in the co-occurrence matrix shown on the top.

We therefore asked whether the full distributions could be described by a mixture of statistical components. To this end, we fitted both avalanche size and silent time distributions without a minimum cutoff value using mixtures of lognormal distributions and compared models containing one, two, and three lognormal components (see Materials and Methods). We found that a mixture of two lognormal components provides a substantially better description of the empirical distributions than the single-lognormal model, and the three-lognormal mixture (as it avoids the over-parameterization introduced by allowing three components).

Interestingly, for both avalanche size and silent time, Fig. 2, panel b), the separation point between the two lognormal components corresponds to the crossing point between the inverse relationship observed in the joint probability distribution. Thus, the statistical separation emerging from the mixture-model analysis independently reproduces the boundaries already suggested by the joint organization of the two variables, suggesting the existence of two preferential dynamical regimes governing avalanche dynamics. Motivated by this observation, we used the intersection points between the two lognormal components as thresholds to partition both avalanche size and silent time into two classes. Combining these classifications yields four possible categories corresponding to the four previously identified regions in the joint probability analysis.

For each avalanche, we compute the probability of belonging to one of the two components of the mixture distribution. Repeating this procedure independently for both sizes and silent times, we then construct, for each event, the co-occurrence matrix of the four possible combinations of sizes and pauses (see Materials and Methods).

We then compared the results with those obtained by recomputing the same matrix after reshuffling the order of subsequent pairs’ size/silent time, as previously done for the joint probability analysis. We observed a positive deviation from the null model along the anti-diagonal elements of the 2 *×* 2 matrix as shown in Fig. 2, panel c) (top). In other words, one quantity (small/large avalanche) co-occurs with the second component (long/short intervals) of the other more frequently than expected under the hypothesis of independence. These results are consistent with those obtained from the joint distribution analysis Fig. 2, panel c) (bottom).

### 3.3 Category persistence reflects the stability of brain dynamics

The results obtained from the joint probability analysis and from the lognormal mixture decomposition lead to a non-trivial labeling of avalanches into four non-overlapping classes (Fig.2, panel c). Using the intersection point of the respective components of the mixture distributions, we identify the four possible combinations of large and small values for avalanche size and silent time. By examining the burst length of consecutive avalanches belonging to the same category, we find that their occurrence is higher than what is observed when reshuffling the temporal sequence of categories while preserving the size–silent time relationship (Fig.3, panel a). We thus observe that a given category can persist over multiple consecutive avalanches more frequently than expected under random dynamics. To explain category persistence in terms of feature dynamics, we estimated the temporal distribution of amplification and attenuation, defined as increases or decreases in avalanche size relative to the preceding event. By modeling the sequences of amplification and attenuation events as Markov chains, with 1 and *−*1 denoting amplification and attenuation, respectively, for both avalanche size and silent time, we found that the probability of remaining in the same state (0.33) was lower than the probability of transitioning to the opposite state (0.66). The non-triviality of this result was assessed by comparison with reshuffled sequences in which the temporal dependence between amplification and attenuation events was destroyed while their relative frequencies were preserved.

**Figure 3:**
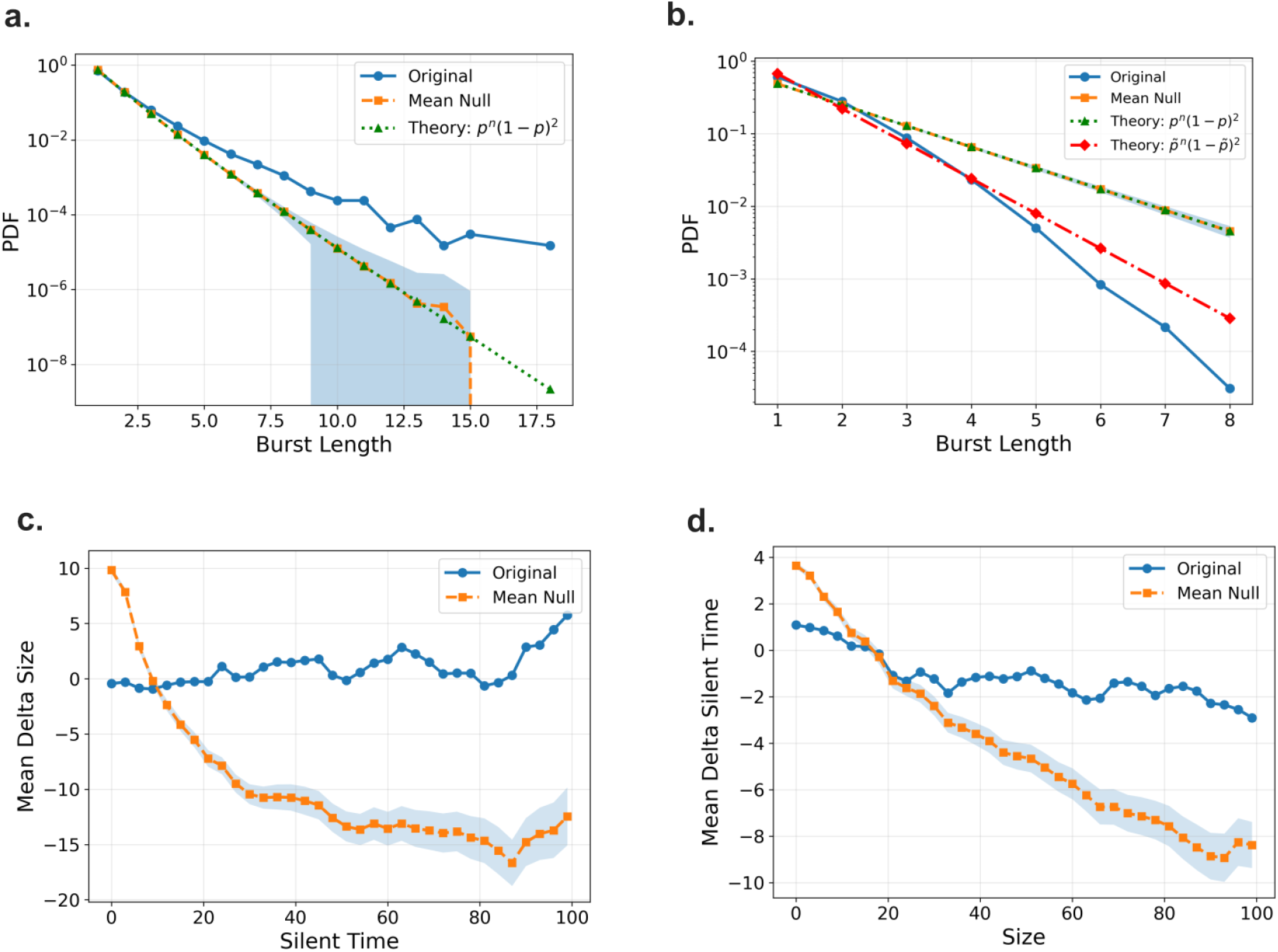
Size–silent time category subdivision and transitions. Panel (a): Probability density function (PDF) of bursts consisting of consecutive events belonging to the same category, compared with a model of independent event occurrences (green line). The model describes the probability of observing *n* consecutive events from a given category when each event is independently assigned to that category with probability *p*.Panel (b): PDF of bursts of consecutive events exhibiting size attenuation, compared with two independent models based on the same principle described in panel (a). Here, *p* represents the probability that the size of an event is smaller than that of the preceding event. The first model (green line) assumes *p* = 0.5, whereas the second model (red line) assumes *p̃* = 0.33. Panel (c): Mean size difference between two consecutive events as a function of the silent time separating them in the original event process (blue curve), compared with the independent event model (orange curve). Panel (d): Mean silent-time difference between two consecutive events as a function of the size separating them in the original event process (blue curve), compared with the independent event model (orange curve).

This effect can be visualized by considering the probability of observing sequences of consecutive amplification or attenuation events (Fig. 3, panel b). The estimated distribution lies below that obtained from reshuffled data, which represents the independent occurrence of the two phenomena.

Consistent with this tendency to alternate between states, sequences of consecutive amplification or attenuation events occurred less frequently than expected from reshuffled data (Fig. 3, panel b). Since reshuffling destroys temporal dependencies while preserving the relative frequencies of the two phenomena, this comparison confirms the non-trivial temporal organization of amplification and attenuation events. This temporal organization is also reflected in the stability of the mean size variation across two consecutive avalanches as a function of the silent time separating them, as well as in the mean silent-time variation conditioned on avalanche size (Fig. 3, panels c and d). In both cases, the observed behavior differs from that of the null model obtained by reshuffling the events while preserving the joint distribution of avalanche sizes and silent times.

The crossing points between the empirical curves and the null model approximately coincide with the intersection points identified in Fig. 2, panel b). Overall, these results indicate that, although the joint probability distribution of avalanche sizes and silent times favors either attenuation or amplification depending on the value of the complementary variable (i.e., silent time when conditioning on avalanche size, and avalanche size when conditioning on silent time), the temporal organization of avalanches tends to balance the occurrence of these two phenomena. This balance mechanism stabilizes avalanche features in consecutive events, thereby contributing to the observed persistence of avalanche categories.

### 3.4 Structural organization of the neuronal avalanches

Resting-state whole-brain activity reflects a structured network organization shaped by the anatomical connections between brain regions. Accordingly, in neuronal avalanches detected from MEG recordings, structural connectivity constrains the propagation of supra-threshold activity across the brain [29]. We therefore hypothesized that structural connectivity influences the properties (i.e., the size) of the avalanches in which each brain region preferentially participates. To investigate this hypothesis, we quantified the propensity of each brain region to participate in small versus large avalanche events (as defined by the partition above. Specifically, for each avalanche-size category, we computed the relative activation frequency (RAF) of each region by counting the avalanches in which that region was active and weighting each occurrence by the inverse of the total number of regions active in the same avalanche. This weighting accounts for the fact that an individual region’s contribution is proportionally smaller in avalanches involving more regions. We found that the RAF varies across brain areas (Fig. 4, Panel a). In particular, some regions exhibit a relative activation frequency greater than 0.5 4, Panel c), indicating a preferential involvement in small avalanches, whereas others show values lower than 0.5, suggesting a stronger engagement during large avalanche events.

**Figure 4:**
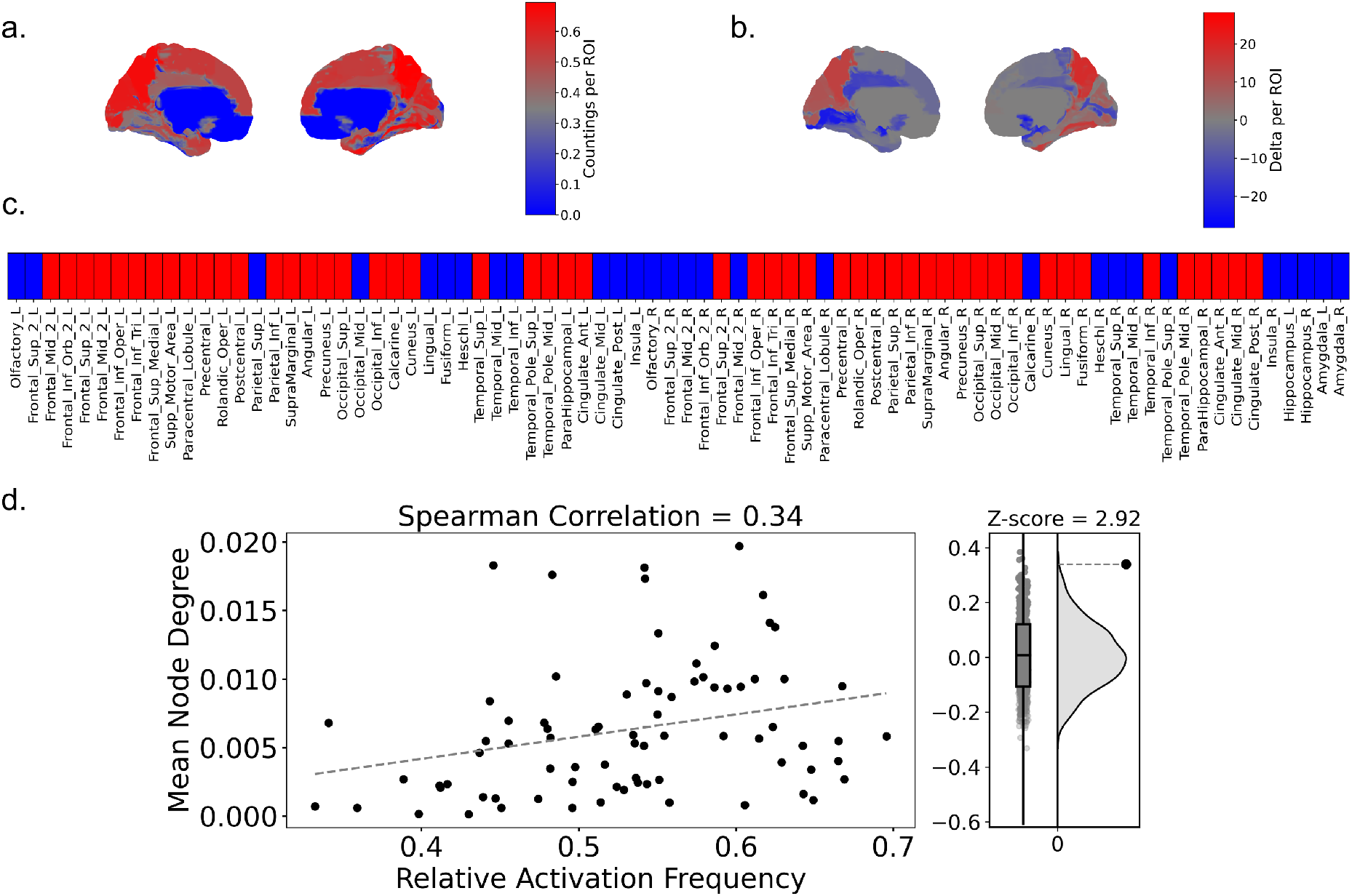
Region activation in small-size events. a) Brain maps showing regional RAF during small-size avalanches. (b) Brain maps showing the z-scored difference between the observed number of regional activations in small-size events and the corresponding null model expectation. (c) AAL atlas regions colored according to RAF during small-size avalanches. Regions with RAF *>* 0.5 are shown in red, whereas all other regions are shown in blue. (d) Spearman correlation between regional relative activation frequency during small-size avalanches and structural node degree (left). The right panel shows the distribution of correlations obtained under the null model, compared with the empirical correlation value.

To assess whether these findings could be explained by random fluctuations, we performed a reshuffling analysis in which active regions within avalanches were randomized while preserving both the avalanche size distribution and the number of regional activations. Under this null model, the observed RAF deviates significantly from chance level (Fig. 4, Panel b), indicating that the observed patterns are not expected by a random organization.

These differences in regional activation appear to be related to the underlying connectivity structure. Indeed, by computing the mean node degree of the structural connectivity for each brain region, we found a correlation (quantified using Spearman’s correlation) with the relative activation frequency (Fig. 4, Panel d). Overall, these results suggest that highly connected regions are most often engaged during small avalanche events. Conversely, the more peripheral a region, the more it is selectively recruited by large bursts.

## 4 Discussion

Neuronal avalanches are transient spatiotemporal cascades of neural activity in which successive neuronal populations are recruited within a temporally contiguous event, bounded by periods of relative quiescence. Their size, duration, and propagation patterns provide a framework for investigating how local fluctuations spread through neural networks and give rise to coordinated activities across spatial scales.

In this work, we characterized whole-brain resting-state dynamics from MEG recordings acquired in healthy subjects. Within the neuronal avalanche framework, we went beyond classical analyses that focus on scaling behaviors, and set out to directly characterize avalanche dynamics. To this end, we introduced a novel approach to assess the relationship between avalanche size and the duration of the silent time preceding avalanche onset, under the hypothesis that the duration of quiescent intervals and the magnitude of the burst are related.

Compared with a null model obtained by randomly reshuffling the correspondence between avalanche sizes and preceding silent times, we found that the empirical joint probability distribution deviates significantly from statistical independence. In particular, small avalanches tend to be preceded by longer silent times, whereas larger avalanches are more likely to follow shorter silent intervals.

This result was further supported by the analysis of the marginal distributions of avalanche size and silent time. Both distributions were well described by a mixture of two lognormal components, suggesting the presence of two distinct dynamical regimes. Notably, in both distributions, the two components intersected near the point at which the empirical joint probability shifted from an excess to a deficit relative to the null model, or vice versa. Thus, the boundaries separating the two marginal regimes also marked a reversal in the coupling between avalanche size and silent time.

To further investigate this relationship, we analyzed the interaction between the two components identified in the mixture models of both avalanche sizes and silent times. Specifically, we computed the probability that an avalanche, characterized by its size and preceding silent time, belongs to each combination of the corresponding mixture components. These probabilities were compared with those expected under a null model assuming independence between components of the avalanche size distribution and components in the silent time distribution. The results confirmed the patterns observed in the joint probability analysis. In particular, the component associated with a higher prevalence of small avalanches is preferentially coupled with the component associated with longer silent periods, whereas the component associated with larger avalanches is more strongly linked to shorter silent intervals.

The presence of these non-random associations further supports the interpretation of the observed distributions as two distinct dynamical regimes.

These findings suggest that the cross-talk between two distinct dynamical mechanisms may shape whole-brain resting-state activity. The observation that both avalanche size and silent-time distributions are better described by mixtures of two lognormal components points toward the coexistence of two statistical regimes rather than a single homogeneous process.

The presence of lognormal mixtures across the full distributions does not necessarily contradict the hypothesis that brain activity operates near a critical point. The mixture model provides a statistical description over the entire range of observed values, whereas scale-free properties may arises only within specific regimes of the dynamics. Finite-size effects and the subsampling inherent to neural recordings can substantially distort the statistical signatures of scale-free processes. In particular, the limited sampling of the underlying system, unavoidable in both microscopic and macroscopic recordings, may lead to deviations from ideal power-law behavior even when the underlying dynamics are critical.

The truncated power law fit above a minumun cutoff revealed a scale-free behavior, with the avalanche-size exponent close to the value of 1.5 predicted by mean-field theories of self-organized criticality.

The origin of the regime characterized by smaller avalanches and longer silent periods does not lend itself to a straightforward dynamical interpretation. Because finite-size effects and subsampling can substantially distort avalanche statistics, the present data cannot establish whether this regime represents another expression of the same critical dynamics or a genuinely distinct operating state of the brain.

Addressing this question will require developing theoretical models that reproduce the observed phenomenology and clarify the mechanisms underlying the coexistence and interaction between the two regimes.

The coexistence of these dynamical regimes is also consistent with results from persistence analyses of dynamical features. Previous studies have reported deviations from independence between avalanche size and silent time [17–20]; however, these approaches typically focused on single-event statistics and did not explicitly address how such features are organized over time across sequences of avalanches.

Here, we extend this perspective by leveraging the intersection point of the mixture distributions to partition the joint feature space into four distinct categories. By assigning each avalanche to one of these categories, we analyzed the temporal organization of the resulting sequence. We found that the same category tends to persist over consecutive events more frequently than expected by chance.

In addition, transitions between categories are not random but tend to alternate between regimes of amplification and attenuation. This suggests that the system remains in a given dynamical state for longer periods than expected by chance. Consistently, although the relationship between avalanche size and silent time associates amplification and attenuation with short and long silent time intervals, respectively, as previously shown in [21], our approach reveals that the temporal organization of avalanches counterbalances this tendency. As a result, positive and negative changes in avalanche features tend to balance each other, producing mean fluctuations close to zero. These fluctuations are *smaller* in magnitude than those predicted by a null model that preserves the joint probability distribution of avalanche sizes and silent times. Overall, these results indicate that the interaction between avalanche sizes and silent times is structured not only at the level of individual events but also across sequences of events, revealing a temporal organization that may reflect the nested timescales at which the brain operates.

The observed persistence of avalanche categories indicates that consecutive neuronal avalanches are not independent events. If avalanches were generated as independent cascades, as commonly assumed in branching-process descriptions of avalanche dynamics, the probability of observing a given category would not depend on the category of the preceding avalanche. Instead, we find that large avalanches preceded by short silent intervals tend to be followed by avalanches with similar characteristics, while small avalanches associated with long silent intervals likewise exhibit enhanced persistence. These findings reveal temporal correlations spanning multiple avalanche events and provide evidence of dynamical memory in the underlying network activity [35], [36]. Such persistence suggests that neuronal avalanches are embedded within longer-lasting dynamical states of the underlying network [25], [26], [37], [38], [39]. Rather than representing isolated cascades of activity, individual avalanches may constitute local manifestations of a global brain state that constrains both activity propagation and extinction. In this framework, sequences of large avalanches separated by short silent times can be interpreted as periods of sustained network activation, whereas sequences of small avalanches interspersed with long quiescent intervals may reflect periods of reduced excitability.

This interpretation is consistent with previous observations of spontaneous alternations between up and down states in neuronal systems [17, 23, 25]. During up states, recurrent network interactions support the propagation of activity and favor temporally clustered avalanches, whereas down states are characterized by longer silent periods and reduced propagation. Alternating between integrated (when avalanches can propagate) and segregated (reduced spreading of large activities) states has been linked to cognitive abilities [40], the presence of conscious perception [41], and neuromodulation [42].brain’s spontaneous alternation between these dynamical regimes.

More broadly, these results support the view that neuronal avalanches are not independent events but are temporally organized by the underlying network dynamics. The observed memory across consecutive events may arise from the persistence of large-scale activity patterns, consistent with theoretical models in which avalanches represent fragments of ongoing attractor-like or replay-like dynamics [25–27] rather than isolated cascades of neural activity.

The identification of multiple dynamical regimes naturally raises the question of whether specific brain regions preferentially contribute to one regime over the other. To address this, we quantified the probability of regional involvement during avalanches, classified as small or large based on the separation derived from the mixture model analysis.

In addition, we examined whether these differences in regional recruitment are related to the brain’s underlying structural organization. In particular, we assessed the relationship between the probability of activation in small avalanches and the mean structural connectivity of each region, estimated as the average node degree in the white-matter connectome. The analysis revealed a nontrivial association: highly connected regions are more frequently engaged during small avalanches, whereas regions with lower connectivity tend to be recruited more prominently during larger events.

Although initially counterintuitive, one possible explanation is that hub regions receive convergent input from many neighboring areas and can therefore be recruited even during relatively weak or spatially confined activity episodes. In addition, highly connected regions may act as preferential sites for the initiation or early recruitment of avalanche activity. In systems operating near criticality, many activity cascades remain spatially limited and terminate before reaching more peripheral regions. As a consequence, hub regions may participate in a large fraction of small avalanches, whereas the recruitment of less connected regions may require stronger and more widespread propagation. Within this perspective, small avalanches preferentially engage the structurally central backbone of the network, whereas large avalanches reflect the progressive recruitment of more peripheral regions.

More generally, the association between avalanche categories and structural connectivity suggests that the dynamical regimes identified here are not purely temporal phenomena but are constrained by the underlying anatomical architecture. This interpretation is compatible with recent whole-brain modelling studies showing that, near criticality, spontaneous activity becomes increasingly shaped by the structural connectome [27]. Within this framework, the persistence of avalanche categories may reflect the system’s tendency to remain transiently confined to specific metastable network configurations, whose expression is constrained by structural connectivity [26, 29, 30]. Overall, our results suggest that neuronal avalanches are organized into persistent dynamical regimes with both temporal and spatial signatures. The persistence of avalanche categories reveals a form of dynamical memory that extends across multiple events, while their spatial organization indicates that these regimes are constrained by the brain’s structural architecture.

In conclusion, these findings suggest that the the dynamical regimes identified through the temporal organization of neuronal avalanches also exhibit distinct spatial signatures, with regional participation being shaped by the constraints imposed by the brain’s structural scaffolding.

## 5 Methods

### 5.1 Data Acquisition

#### Participants

Thirty young healthy adults (mean age 27.28 *±* 6.78) were enrolled in the study. All participants were right-handed and native Italian speakers. Eligibility criteria required: (1) no history of relevant internal, neurological, or psychiatric disorders; and (2) no use of medications or substances potentially interfering with MEG or MRI measurements. The study was conducted in accordance with the Declaration of Helsinki and was approved by the local Ethics Committee. All participants provided written informed consent prior to participation.

#### MRI acquisition

Structural three-dimensional T1-weighted images were acquired using a 1.5 T MRI scanner (Signa, GE Healthcare) and a 3D Magnetization-Prepared Gradient-Echo BRAVO sequence. Acquisition parameters were TR/TE/TI = 8.2/3.1/450 ms, isotropic voxel size of 1 mm^3^, 50% overlapping partitions, and 324 sagittal slices covering the entire brain. Diffusion-weighted MRI data were additionally acquired to reconstruct subject-specific structural connectomes. An Echo-Planar Imaging sequence was used with TR/TE = 12,000/95.5 ms and a voxel resolution of 0.94*×*0.94*×*2.5 mm^3^. Diffusion weighting was applied along 32 directions, including five B0 volumes. MRI acquisition was performed after the MEG recording session. Diffusion MRI data were preprocessed using tools from the FMRIB Software Library (FSL, http://fsl.fmrib.ox.ac.uk/fsl/). Head-motion and eddy-current distortions were corrected using the *eddy-correct* routine, including the appropriate reorientation of the diffusion-sensitizing gradient directions. Brain masks were subsequently obtained from the B0 images using the Brain Extraction Tool. A diffusion-tensor model was estimated voxel-wise, and whole-brain streamlines were reconstructed through deterministic tractography using the Diffusion Toolkit. Fiber tracking was performed with the FACT algorithm, adopting a 45*^◦^* angular threshold and spline filtering. Fractional Anisotropy (FA) maps, thresholded at 0.2, were used as tracking masks. For the tractography analysis, ROIs from the AAL atlas and a volumetric implementation of the DKT atlas were considered. Both atlases were defined in MNI space and restricted using the gray-matter (GM) probability map obtained with SPM, applying a threshold of 0.2. For each participant, FA volumes were registered to MNI space using the FSL FA template together with the spatial normalization procedure implemented in SPM12. The resulting transformations were inverted and applied to the ROIs to map the parcellations into each participant’s individual space. The accuracy of the normalization procedure was visually inspected. Finally, subject-specific whole-brain tractography and the corresponding GM ROI sets were processed using custom software developed in Interactive Data Language (IDL; Harris Geospatial Solutions, Inc.). For each pair of GM ROIs, structural connectivity was quantified by computing the number of connecting streamlines and their mean tract length.

#### MEG pre-processing

MEG recordings were obtained using a system developed by the National Research Council of Italy at the Institute of Applied Sciences and Intelligent Systems (ISASI), comprising 163 magnetometers [43]. Resting-state activity was recorded with participants keeping their eyes closed and was divided into two runs of 3 min and 30s each. Head position was estimated using four anatomical landmarks and four position coils, whose locations were digitized. Electrocardiographic (ECG) and electro-oculographic (EOG) signals were simultaneously recorded to facilitate the identification of physiological artifacts. MEG signals were acquired at a sampling frequency of 1024 Hz after application of an anti-aliasing filter. The recordings were subsequently band-pass filtered between 0.5 and 48 Hz using a fourth-order Butterworth IIR filter. Environmental interference measured by the reference magnetometers was attenuated through principal component analysis. Physiological artifacts, including ocular and cardiac components, were identified and removed using supervised independent component analysis, typically resulting in the exclusion of one component. Channels exhibiting excessive noise were manually identified and discarded by an expert rater. Following this quality-control procedure, an average of 136 *±* 4 sensors per participant were retained for subsequent analyses.

#### Source reconstruction

Source-level neuronal time series were initially reconstructed for 116 regions of interest (ROIs) defined according to the AAL atlas. The forward model was obtained using the volume-conduction approach introduced by Nolte [44], based on each participant’s individual structural MRI. Source activity was then estimated using a linearly constrained minimum variance (LCMV) beamformer [45], with reconstruction performed at the centroid of each ROI. For the analyses presented here, the original set of 116 ROIs was reduced to 78 cortical regions to restrict the analysis to the most reliable signals. All MEG preprocessing and source-reconstruction procedures were carried out using the FieldTrip software package [46].

### 5.2 Avalanches Detection

Avalanches in neuronal systems are defined as bursts of coordinated activations across multiple regions [1]. While in spiking data the discretization of the signal into spiking versus non-spiking activity naturally provides a definition of regional activation, in continuous signals such as MEG recordings determining whether a region is active requires a specific analysis to assess how much the signal exceeds a defined baseline level of activity [5]. In doing so, we can define a threshold above which the signal amplitude is considered indicative of regional activation. The signal can then be binarized, classifying each region as either active or inactive depending on whether its activity exceeds the selected threshold. To define this threshold, we z-scored the signal for each region. We then computed the grand-average distribution of the z-scored signal across all recorded regions and all the subjects. By fitting the obtained distribution, we compared it with a Gaussian distribution having the same mean and standard deviation as the estimated distribution. We observed that for values greater than 3 and, symmetrically, for values smaller than *−*3, the empirical distribution began to deviate substantially from the corresponding Gaussian distribution. Therefore, we binarized the region-by-time matrix by assigning 1 to entries with z-scored signal exceeding 3 and 0 otherwise. To reduce the influence of noisy processes, we applied temporal binning to the binarized matrix, where each new bin was obtained by summing 3 consecutive temporal bins. To identify avalanches in the binarized matrix, we applied the procedure used in [5]. Starting from a non-avalanche state (i.e., a time bin in which no regions are active), an avalanche is considered to begin at time *t* when at least one region becomes active. The avalanche continues as long as there is no time bin in which all regions are inactive (i.e., all entries equal to zero). During this period, the active regions at each time bin are recorded. The avalanche terminates when a time bin has no active regions, and the system remains in this non-avalanche state until a new activation event initiates the next avalanche. Once the entire signal had been segmented into avalanches, we computed several metrics for each event. The duration was defined as the number of temporal bins during which the avalanche remained active. The size was calculated as the sum of active bins across all active regions, multiplied by the duration of the avalanche. The active regions were defined as those that were active at least once during the avalanche. Finally, the silent time was defined as the number of temporal bins between the end of one avalanche and the onset of the subsequent one. The features extracted within the avalanche framework were then concatenated across subjects.

### 5.3 Joint Probability distribution

The joint probability distribution of avalanche size and preceding silent time was estimated using logarithmically spaced bins for both variables. For each bin, we computed the empirical probability of observing a silent time–size pair within the recorded time series. To assess deviations from statistical independence between avalanche size and preceding silent time, we constructed a null model in which avalanche sizes were randomly reshuffled while preserving the temporal sequence of silent times. Thus, each silent interval was reassigned a randomly selected avalanche size, leaving the marginal distributions unchanged while destroying any potential dependence structure. For each reshuffling, the joint probability distribution was recomputed. Since reshufflings were performed independently, the null distribution of probabilities within each bin was approximately Gaussian, in accordance with the central limit theorem. For each bin, we estimated the mean and standard deviation of the null distribution and quantified deviations of the empirical distribution by computing the corresponding z-score, defined as the standardized difference between the observed probability and the null model mean. We validated this procedure using a bootstrap approach across subjects. Specifically, we randomly selected 20 subjects, concatenated the features extracted from each, and recomputed the joint probability distribution. This random selection was repeated 50 times. We then averaged the z-scored joint distributions across all iterations and compared the resulting mean distribution with that obtained by concatenating features from all subjects, confirming the robustness of the results.

### 5.4 Lognormal Mixture interploation

To determine whether the PDFs of avalanche sizes and silent times are better described by a mixture of two lognormal distributions rather than a single lognormal distribution, we used the Python package GaussianMixture to fit the mixture model and extract its components. We then applied two statistical criteria for model comparison with different numbers of degrees of freedom, namely the Akaike Information Criterion (AIC) [47] and the Bayesian Information Criterion (BIC) [48]. Both criteria indicated that a mixture of two lognormal distributions provides a better fit to the empirical PDFs. Once the two components of the respective distributions were obtained, we investigated whether and how these components are mutually dependent between avalanche size and silent time. To this aim, we first estimated, for each event, the posterior probability of belonging to each mixture component for both size and silent time. We then quantified the co-occurrence structure of components across the two variables. Within this framework, let 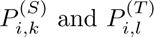 denote the posterior probabilities that event *i* belongs to component *k ∈ {*1, 2*}* of the size distribution and to component *l ∈ {*1, 2*}* of the silent-time distribution, respectively.

We then define the component co-occurrence matrix ***ω*** *∈* R^2*×*2^ as:

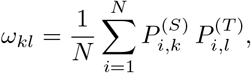

where *N* is the total number of events.

Each element *ω_kl_* represents the expected joint probability that a given event simultaneously belongs to component *k* of the size mixture model and component *l* of the silent-time mixture model, thereby providing a measure of coupling between mixture components across the two variables. We then compared the results obtained from the empirical data with those of a null model, in which the temporal correspondence between size and silent time was destroyed by randomly reshuffling the ordering of the size and silent time pairs across events, as we did in Subsection 5.3.

### 5.5 Category Persistency and Mean Variation

The separation of the lognormal mixtures allowed us to identify a precise threshold between the two putative mechanisms governing avalanche size and silent time dynamics. By combining the respective components associated with the two quantities, we assigned a numerical label to each possible combination, thereby defining the event categories shown in panel c of Fig. 2. In this way, we obtained a vector describing the temporal succession of event categories.

We then computed the probability of observing bursts of consecutive events belonging to the same category as a function of burst size. The results obtained were compared with those from a null model generated by reshuffling the category sequences. In this case, the reshuffling procedure was performed by applying the same permutation index simultaneously to both avalanche sizes and silent times. This procedure destroys the temporal ordering of events while preserving the marginal distributions of the two quantities and their mutual dependence, as encoded in the joint probability distribution.

The probability of observing bursts of identical categories in the original data was consistently higher than that obtained from the reshuffled sequences for each category individually (see Supplementary Materials), as well as when considering all categories together, as shown in Fig. 3. Furthermore, the null distribution was compared with a model based on random and independent extraction of categories, confirming that the persistence of a given category in the original series is significantly higher than expected from independent occurrences.

To further support this result, we computed the sign of the size variation for each pair of consecutive events, yielding a binary sequence of *−*1 and +1 values corresponding to attenuation and amplification states, respectively. We first estimated the associated Markov chain transition matrix to quantify the probability of transitions between attenuation and amplification states. We found that the transition probability between the two states (*∼* 0.66) was higher than the probability of remaining in the same state.

Using this information, we computed the burst persistence distributions for the attenuation and amplification states, following the same procedure as in the category analysis. In this case, the empirical results were compared with a null model obtained by randomly reshuffling the sign sequence. This reshuffling procedure preserves the total number of attenuation and amplification events while removing their temporal ordering.

Finally, we computed the mean variation of avalanche size and silent time as a function of the value intervals of the complementary variable. More specifically, we evaluated the average size variation conditioned on silent time intervals and, conversely, the average silent time variation conditioned on avalanche-size intervals.

To assess the significance of the observed relationships, the results were compared with those from a null model generated by jointly reshuffling the size and silent-time arrays with the same permutation index.

### 5.6 Regions activations in small avalanche events

To assess whether the structural connectome biases regional recruitment toward one of the two avalanche-size categories (below or above 20), we quantified, for each region, the relative proportion of activations in each category. Since avalanche size is, by definition, proportional to the number of active regions, larger avalanches inherently involve more regions. To avoid a trivial inflation of activation probability in the large-size category due solely to the greater number of recruited regions, we weighted each activation by the inverse of the total number of active regions within the corresponding avalanche. In this way, each avalanche contributed equally, independently of its size, preventing large avalanches from disproportionately influencing the estimated regional probabilities. Once the proportional activation counts for each region in the two size categories were obtained, we tested the non-triviality of these results using a null model. Specifically, we reshuffled the assignment of avalanches to the two size categories while keeping the total number of avalanches in each category and the specific pattern of active regions within each avalanche fixed. In this way, the internal structure of each avalanche was preserved, while its membership in one of the two size categories was randomized. This procedure tests the null hypothesis that regional recruitment is homogeneous with respect to avalanche size, i.e., each region has the same probability of appearing in avalanches of either category. For each reshuffling, we recomputed the proportional activation counts, thereby obtaining a null distribution for every region. We then quantified the deviation of the empirical value from the null expectation by computing the standardized score (z-score), defined as the difference between the observed value and the mean of the null distribution, normalized by the null distribution’s standard deviation. Finally, we computed the Spearman rank correlation between the regional proportion of activations and the mean node degree derived from the structural connectivity matrix. To assess the non-triviality of the observed correlation, we constructed a second null model in which the mean node degrees were randomly reshuffled across regions, thereby preserving the overall degree distribution while breaking the specific association between structural connectivity and regional activation probability. The empirical correlation coefficient was then compared with the corresponding null distribution, and statistical significance was quantified using the z-scored distance between the observed correlation and the mean of the null model.

## Acknowledgments

This work was supported by the Governo Italiano, Ministero per lo Sviluppo Economico, ACCORDI PER INNOVAZIONE, “Approccio User-friendly integrato per Diagnosi, Assistenza e Cura Efficaci” (AUDACE), grant number B69J23006050007.

This work was also supported by the Italian Research Infrastructure for Neuroscience (IRIN), funded by NextGenerationEU/Italian National Recovery and Resilience Plan (NRRP), M4C2 (Project code IR0000011, CUP B51E22000150006).

## Code availability

The analysis code and Jupyter notebooks used in this study are publicly available at https://github.com/GianmarCafaro/Avalanche-and-temporal-Persistency-for-MEG-data. The repository includes documentation and instructions for reproducing the analyses. The research data are not publicly available.

